# Simultaneous and precise generation of *Zebra3 and Wsl5* mutations in rice using CRISPR/Cas9-mediated adenine base editors

**DOI:** 10.1101/784348

**Authors:** Kutubuddin A. Molla, Justin Shih, Yinong Yang

## Abstract

The CRISPR/Cas9-mediated base editing technology can efficiently generate point mutations in the genome without introducing double-strand break (DSB) or supplying a DNA donor template for homology-dependent repair (HDR). In this study, adenine base editors (ABEs) were used for rapid generation of precise point mutations in two distinct genes, *OsWsl5*, and *OsZebra3 (Z3)*, in rice protoplasts and regenerated plants. The precisely engineered point mutations were stably inherited to subsequent generations. These single nucleotide alterations resulted in single amino acid changes and associated *wsl5* and *z3* phenotypes as evidenced by white stripe leaf and light green/dark green leaf pattern, respectively. Through selfing and segregation, transgene-free, base edited *wsl5* and *z3* mutants were readily obtained in a short period of time. We noticed a novel mutation (V540A) in *Z3* locus could mimic the phenotype of *Z3* mutation (S542P). Furthermore, we observed unexpected non-A/G or T/C mutations in the ABE editing window in a few of the edited plants. The ABE vectors and the method from in this study could be used to simultaneously generate point mutations in multiple genes in a single transformation and serve as a useful base editing tool for crop improvement as well as basic studies in plant biology.

**Highlights:** Adenine base editors were adapted for plant base editing that can generate precise and heritable point mutations in rice genome without indel formation. The base editing approach allows rapid generation of transgene-free rice mutants with expected phenotypic changes.

## Introduction

Precision genome editing is a powerful tool for accelerating crop improvement. The discovery of the CRISPR/Cas9 system and its repurposing for genome editing revolutionized basic biological research and practical applications in medicine and agriculture (Doudna and Charpentier, 2014). In the CRISPR/Cas9-mediated genome editing process, a single guide RNA (sgRNA) can direct Cas9 to create a double strand break (DSB) at a target locus with high precision. Higher eukaryotes including plant cell repairs the DSB through either non-homologous end joining (NHEJ) or homology directed repair (HDR) pathways (Molla and Yang, 2019*a*). Higher plants predominantly use the error prone NHEJ which creates random insertion/deletion (indel) causing frameshift mutation and ultimately gene knock out (Huang and Puchta, 2019).

NHEJ-mediated gene knock-out has limited application in crop improvement since it cannot create precise indels or specific point mutations for sophisticated crop genome engineering. To install precise point mutations and to delete or introduce desired DNA sequences, we highly rely on HDR-mediated precise genome editing. For HDR to occur, one needs to supply an exogenous donor template containing the desired changes flanked by homologous sequences to the target locus. Unfortunately, the efficiency of HDR is extremely low in higher plants because of the low innate rate of cellular HDR and difficulties in donor template delivery (Huang and Puchta, 2019).

The ability to efficiently generate point mutations in plants has great potential to assist in crop improvement as well as to unravel functions of many natural single nucleotide polymorphisms (Molla and Yang, 2019*b*). CRISPR/Cas-mediated base editing systems have recently been developed to precisely install point mutations in a genome (Komor *et al.*, 2016; Nishida *et al.*, 2016; Gaudelli *et al.*, 2017). Using cytidine deaminase fused with dCas9 (catalytically dead) or nCas9 (nickase), precise C>T base conversions were achieved in both mammalian and plant systems (Komor *et al.*, 2016; Nishida *et al.*, 2016; Shimatani *et al.*, 2017; Zong *et al.*, 2017). Fusion of a hypothetical DNA adenine deaminase with dCas9/nCas9 would generate an adenine base editor, but all such naturally occurring enzymes deaminate adenine only in RNA substrates (Gaudelli *et al.*, 2017)^4^. In order to develop an A>G (and T>C) base editing system, the *E. coli* tRNA adenosine deaminase (TadA) has been modified to accept DNA as a substrate through extensive directed evolution (Gaudelli *et al.*, 2017). This laboratory-evolved DNA adenine deaminase, tethered to nCas9, can now deaminate adenine (A) in the non-target strand into inosine (I). Since inosine is read as guanine by cellular replication machinery, the deamination results in a post-replicative conversion of A to G (Alseth *et al.*, 2014; Gaudelli *et al.*, 2017). Among the several adenine base editors (ABE) developed, ABE7.10 is the most efficient when the target A is within the 4-7 protospacer positions, whereas ABE7.9, ABE7.8, and ABE6.3 perform better than ABE7.10 within the 8-10 protospacer positions (Gaudelli *et al.*, 2017). Although a few of studies have recently been published while preparing this manuscript (Hua *et al.*, 2018*a*; Kang *et al.*, 2018; Li *et al.*, 2018; Yan *et al.*, 2018), relatively little is known about ABE’s wide applicability in the plant system and the heritability of the induced mutation in the subsequent generation in rice plants. Moreover, Most of the earlier studies described ABE induced mutation generation without any phenotypic evidence (Hua *et al.*, 2018*a*,*b*; Yan *et al.*, 2018).

To further demonstrate the application of adenine base editing technology in plants, we attempted to simultaneously edit the *White stripe leaf 5* (*Wsl5*) and *Zebra3* (*Z3*) loci in rice (*Oryza sativa* ssp. *japonica* cv. *‘*Kitaake*’*), a representative cereal crop and monocot model. *OsWSL5*, a recently characterized rice gene, encodes a novel chloroplast-targeted pentatricopeptide repeat protein, which plays an essential role in rice chloroplast biogenesis (Liu *et al.*, 2018). A single nucleotide polymorphism (T to C) located in the conserved region of exon 1 causes a leucine to proline amino acid substitution in the *wsl5* mutant, which can be phenotypically visualized as longitudinal albino leaf striations. *OsZ3*, another recently characterized rice gene, encodes a citrate transporter (Kim *et al.*, 2018). The mutant plant possesses a single base substitution (T to C) in the third exon, causing a missense mutation (serine to proline at amino acid 542), with mutant *z3* plants exhibiting a phenotype of alternating transverse dark-green/light-green stripes in the leaves and growth stunting.

Here we report the development of a plant base editing system based on *E. coli* TadA-derived adenine base editors, demonstrate its utility for single nucleotide mutations and their germline transmission, and provide evidence for the associated mutant phenotypes. This system allows to install non-HDR-based indel-free and multiplexed base change mutations, and to readily generate transgene-free, single base edited plants through selfing or backcrossing.

## Results

### Adenine base-editor vectors for targeted base editing in plants

To perform targeted adenine base editing in rice, we have constructed a vector (*pPr-ABE7.10*) for plant adenine base editing for protoplast transient assays, as well as two binary vectors (*pKABE7.10* and *pKABE7.9*) for *Agrobacterium*-mediated plant transformation (Figure 1A). All the vectors have a similar basic structural configuration composed of an engineered adenosine deaminase fused with a Cas9 nickase (nCas9-D10A). To carry out the deamination reactions, a heterodimeric adenosine deaminase composed of a wild type tRNA adenosine deaminase (TadA) and an engineered TadA (TadA*) has been tethered to nCas9 using a linker peptide. The fused TadA-TadA* -nCas9 ORF was constitutively expressed under the control of a rice ubiquitin promoter. The polycistronic tRNA-gRNA (PTG) gene was expressed under the control of a polymerase III promoter, OsU3 (Figure 1B). Both the binary vectors contained hygromycin resistant gene as a selectable marker.

**Figure 1.**
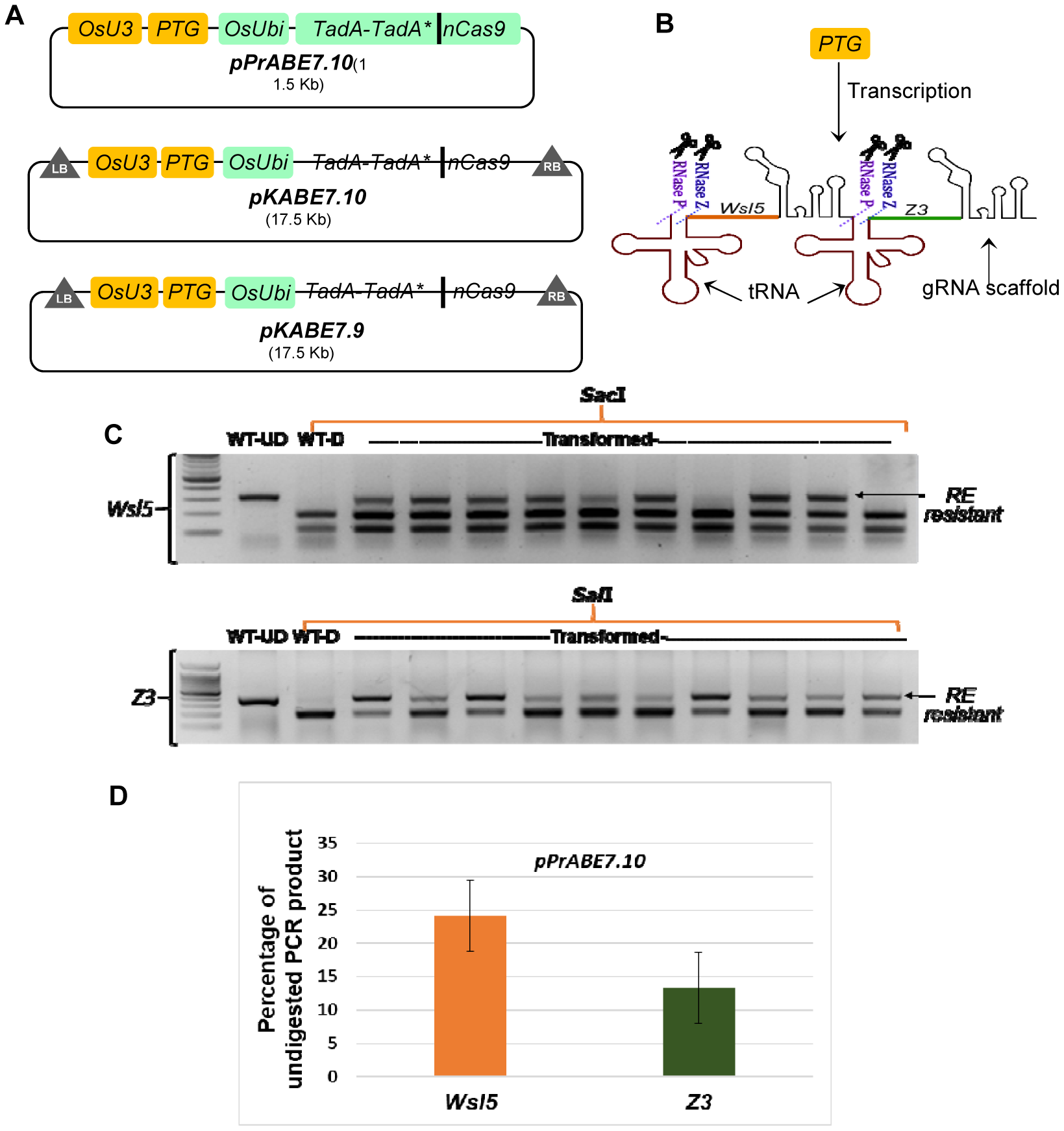
A→G base editing efficiencies at two target loci in transient assays using rice protoplasts. **A.** Schematic diagram of plant adenine base editors constructed and used in this study. *pPrABE7.10* was used in protoplast transformation; *pKABE7.10* and *pKABE7.9* were used for *Agrobacterium*-mediated transformation. **B.** Diagrammatic representation of how PTG (polycistronic tRNA-gRNA) gene works in generating more than one single guide RNA (sgRNA). Endogenous RNaseP and RNaseZ splice out the tRNA releasing sgRNAs for targeting *Wsl5* (orange protospacer) and *Zebra3* (green protospacer) loci. **C.** Representative image of a 2% agarose gel showing a restriction enzyme (RE) digestion of PCR amplicons spanning the target loci. RE resistant band indicates disruption of recognition sites by base editing. *Sac*I and *Sal*I enzymes were used for *Wsl5* and *Z3*, respectively. WT-UD, wild type undigested/untreated; WT-D, wild type digested with RE. **D.** Percentage of undigested PCR product after overnight incubation with RE. Bar diagram represents result for *n*=15±SE.

The original *ABE7.10* and *ABE7.9* base editors were designed for slightly different activity windows (Gaudelli *et al.*, 2017). The *ABE7.10* based vectors are suitable to edit target bases at protospacer position 4-7 (counting the PAM as 21-23), whereas *ABE7.*9 based vectors are better suited for target bases positioned at 7-10. We have cloned two tandemly arrayed guide-RNAs into our PTG, which target the loci *OsWsl5* (*OsKitaake04g315500.1*) and *OsZebra3* (*OsKitaake03g038700.11*). For the *OsWsl5* locus, the target A is at the 5^th^ position of the protospacer, whereas the target A for *OsZ3* locus is at the 7^th^ position of the protospacer.

### Adenine base editing in protoplasts via transient expression assay

To get a rapid indication on whether adenine base editors are functional in rice cells, we transfected Kitaake protoplasts with the *pPr-ABE7.10* vector, containing a PTG for targeting *Wsl5* and *Z3* loci. After 48 hrs of incubation, both the target loci were amplified from the transfected protoplast DNAs by PCR. To detect mutations, we performed cleaved amplified polymorphic sequence (CAPS) analysis using *Sac*1 and *Sal*1 restriction endonucleases (RE) for *Wsl5* and *Z3*, respectively. Successful conversion of the target A to G abolishes a *Sac*1 site (3’-CTCG**A**G-5’) in the *Wsl*5 protospacer. A 310 bp amplified wild type *Wsl5* fragment was completely digested by *Sac*I into 197 and 113 bp fragments, whereas the PCR fragment amplified from transfected protoplast DNA showed partial resistance to cleavage (Figure 1C). Similarly, targeted conversion of A to G in the *Z3* locus destroys a *Sal*I restriction site (3’-C**A**GCTG-5’). Amplified 403 bp wild type *Z3* fragment should be digested into 205 and 188 bp upon *Sal*I treatment. Gel profiles of *Sal*I digested PCR product revealed that targeted mutations were induced in a portion of the protoplast population (Figure 1C). As there were no other A bases in the targetable window, successful mutation of the specific A could be screened by detecting undigested bands (RE resistant bands). Based on the band intensity, we have observed an average of 24.14% and 13.31% RE resistant bands for *Wsl5* and *Z3* loci, respectively (Figure 1D). Editing efficiency was determined by the percentage of PCR amplicons insensitive to restriction digestion. Subsequent sequencing of these PCR amplicons further confirmed successful targeted A>G base editing at both loci in rice protoplasts (Supplementary figure 1).

### Single base conversion at *Wsl5* and *Z3* loci in regenerated plants

Because we observed less base editing efficiency at the *Z3* locus in our protoplast assay (Figure 3), we hypothesized that the lower efficiency might be due to the position of the targeted base. The target A for *Z3* is more proximal (7th) to the PAM than the target base (5th) for *Wsl5*. Therefore, we decided to use both *pKABE7.10* and *pKABE7.9* binary constructs for stable transformation and regeneration of transgenic plants. Based on the earlier report in the mammalian system, ABE7.9 could be more efficient than ABE7.10 for a target A proximal to PAM (Gaudelli *et al.*, 2017). A total of 160 and 165 hygromycin resistant transgenic plants were regenerated with the *pKABE7.10* and *pKABE7.9* binary constructs, respectively. To evaluate base editing, genomic DNA was isolated from each of the regenerated plants and initially screened for the presence of base editors by ABE specific PCR primers. We obtained 142 and 149 ABE-positive plants for *pKABE7.10* and *pKABE7.9*, respectively. From these ABE-positive transgenic plants, the target loci (*Wsl5* and *Z3*) were amplified by PCR. RE analysis of the PCR amplicons was initially used to calculate the editing efficiency. Amplicons from successfully edited plants were found to be partly insensitive to restriction digestion (data not shown). To verify the result, we performed Sanger sequencing and confirmed the single base conversions that are consistent with the RE assay.

**Figure 2.**
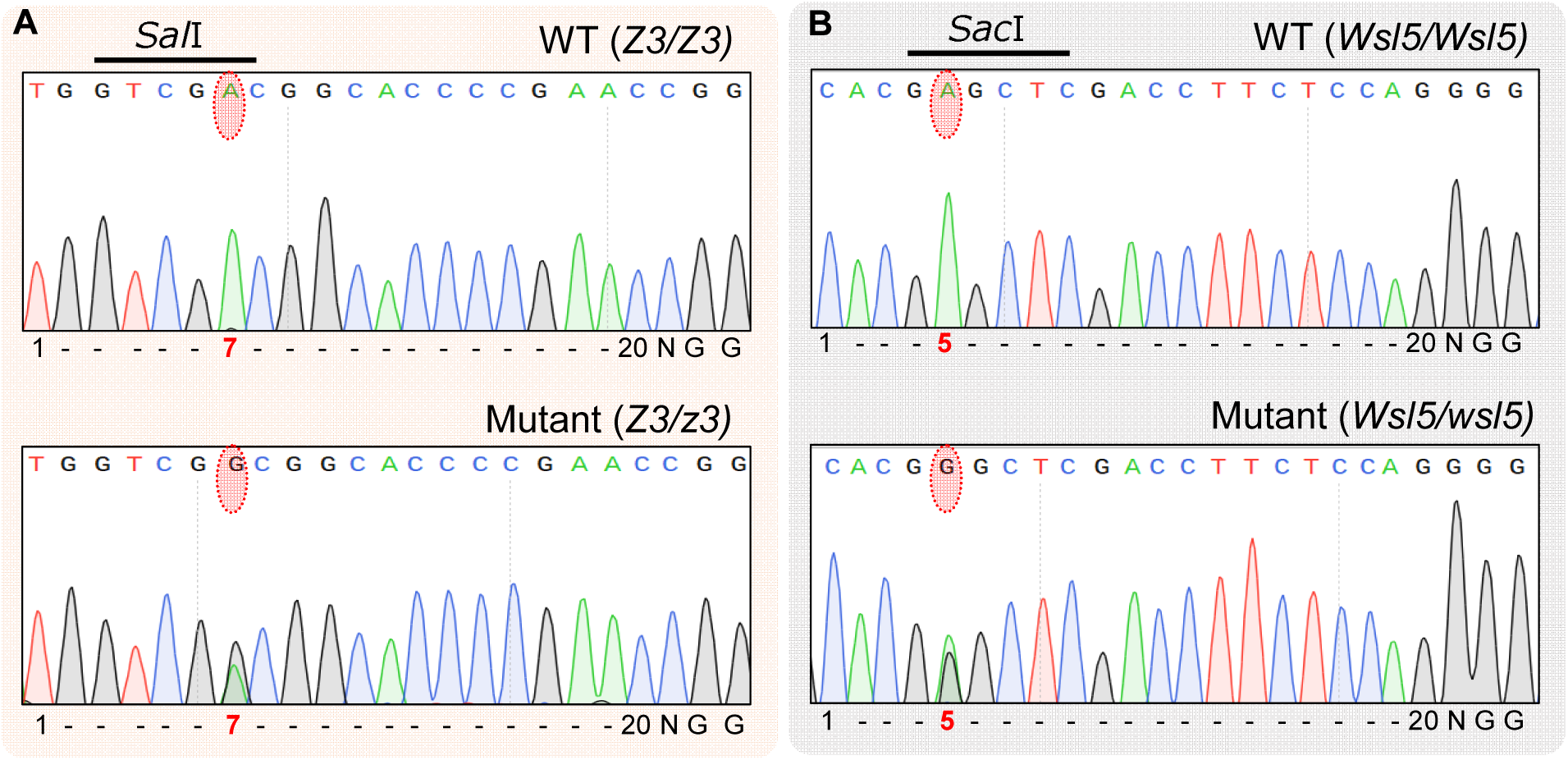
Chromatogram showing successful monoallelic targeted A→G editing in T0 regenerated rice plants. **A.** Editing of 7^th^ A in the *Z3* protospacer, **B.** Editing of 5^th^ A in the *Wsl5* protospacer. Targeted and resultant bases are encircled with red dotted bubble. Upper and lower panels represent chromatogram from wild type and T0 base edited plants, respectively.

**Figure 3.**
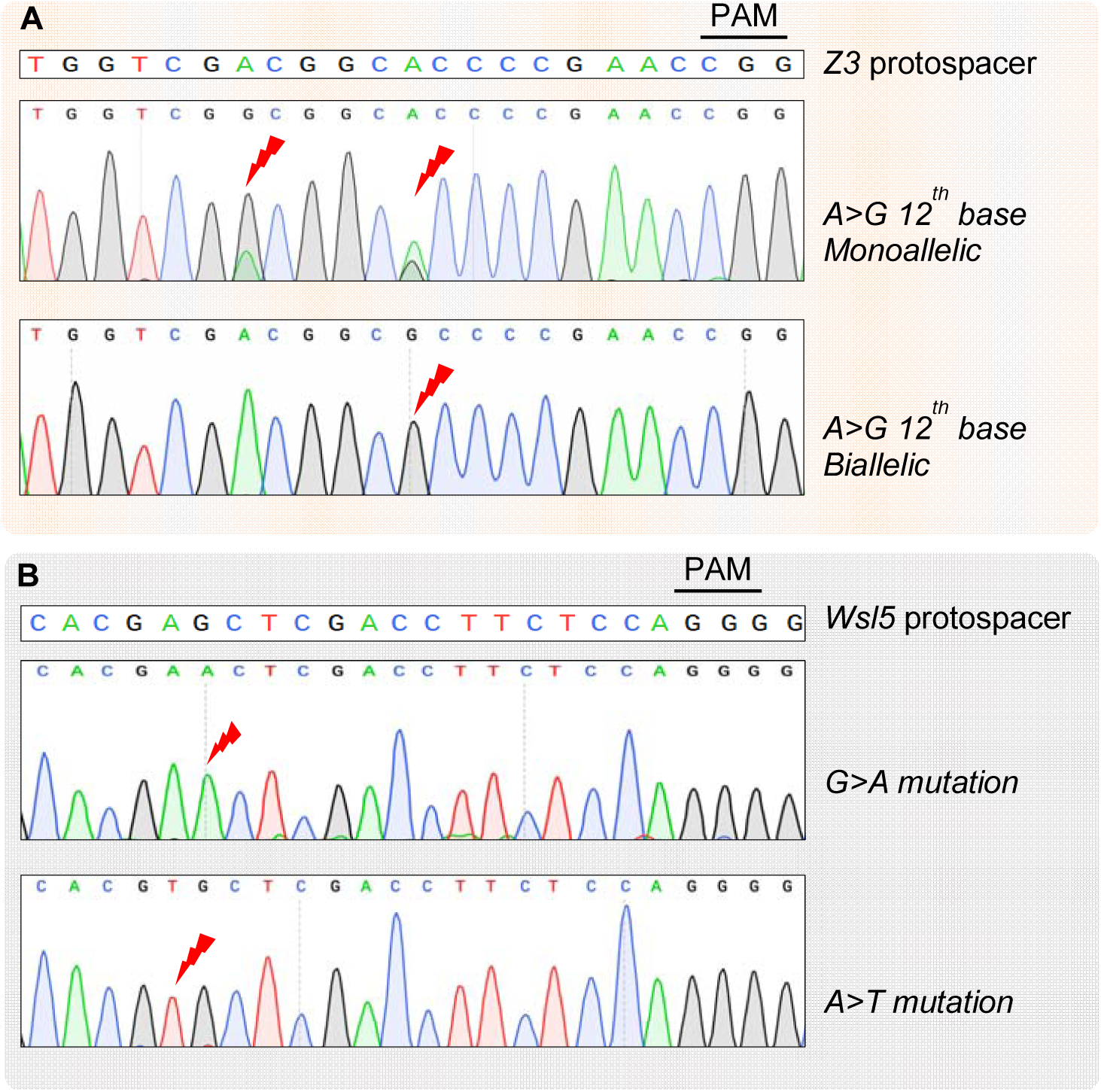
Unintended base editing in T1 plants by adenine base editors. **A.** Monoallelic and biallelic editing of 12^th^ A in the *Z3* protospacer seq. **B.** G→A and A→T conversion in the *Wsl5* protospacer.

Out of the 142 plants derived from *pKABE7.10* transformation, a total of 20 (14.08%) plants were found to be successfully edited for the *Z3* locus, whereas only 4 (2.81%) plants were obtained with the edited *Wsl5* locus (Table 1). On the other hand, *pKABE7.9* showed a much higher editing efficiency (38.92%) for the *Z3* locus. Again, the target base at *Wsl5* locus remained recalcitrant to editing, showing success only in two (1.34%) out of 149 plants (Table 1). Considering both constructs, for *Wsl5*, a total of six plants with intended A to G conversion were obtained exhibiting 1.7% editing efficiency. Interestingly, all the edited T_0_ plants obtained were heterozygous in edited loci as evidenced from partial sensitivity to RE digestion and overlapping peaks in the chromatograms (Figure 2). Analysis of Sanger sequencing data from T_0_ plants does not reveal any unintended proximal base editing or indel formation (Figure 2). Collectively, these experiments demonstrate the feasibility of simultaneous generation of single base edited mutants for two different genes. While we initially intended to obtain individual plants with both sites mutated, due to the low efficiency of base editing at the *Wsl5* locus, we were unable to obtain stable transgenic lines with both types of mutations.

**Table 1.**
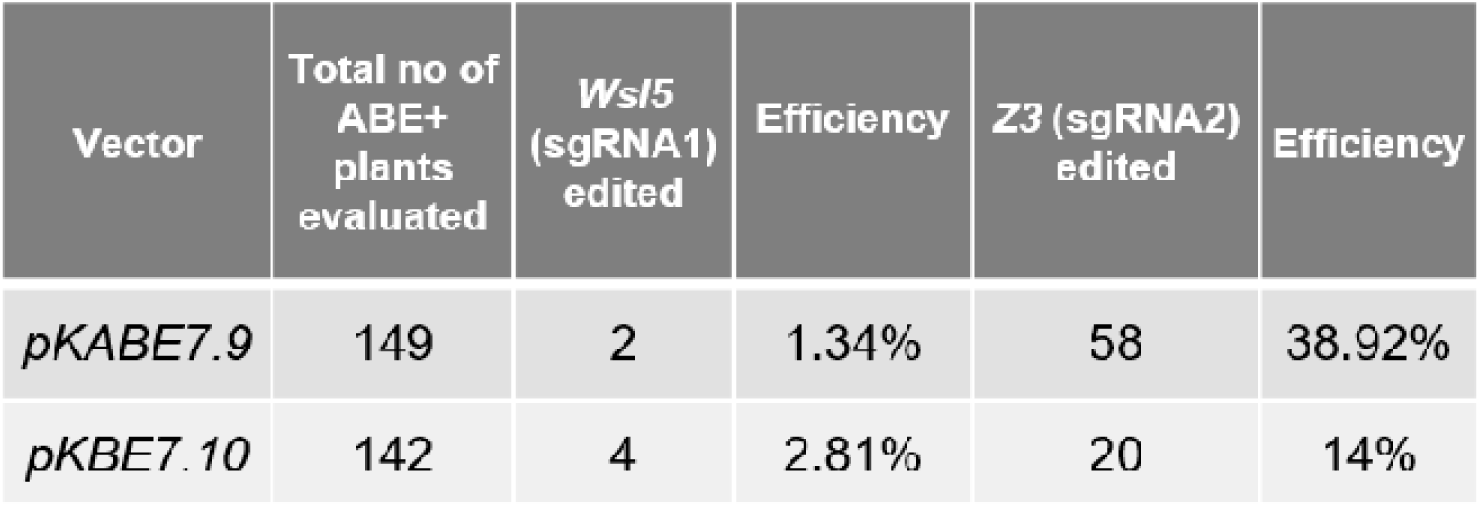
Base editing efficiencies in regenerated rice plants.

### Stable inheritance and segregation of targeted mutations

Next, we sought to follow segregation of the targeted mutations in the T_1_ generation. T_1_ plants derived from the self-pollination of T_0_ monoallelic mutant lines were subjected to inheritance pattern analysis. Like the analysis done for the T_0_ plant, we performed targeted amplification and RE analysis from the T_1_ plants. Among the four *Z3* mutant lines tested, segregation of the mutation in two lines (10-105, and 9-38) did not follow the expected Mendelian ratio (1:2:1). The other two lines (10-51, and 9-86) followed the Mendelian law as evidenced from the calculated chi-square (χ^2^) value being less than the critical value (*p* < 0.05). Similarly, among the *Wsl5* mutant lines, one line (9-79) exhibited deviation from Mendelian segregation pattern and two lines (10-145, and 9-72) showed segregation according to Mendelian law. On the contrary, when we calculated the segregation pattern of base-editors based on ABE specific PCR, all three lines showed inheritance following Mendelian 3:1 ratio. Taken together, these results suggest that the mutation generated by plant adenine base editors are stably inherited to the next generation.

### Unintended proximal base editing in T_1_

Unintended editing in non-target bases of the protospacer and in the nearby region is termed as proximal base editing. Out of 84 mutant T_0_ plants, not a single plant exhibited unintended proximal base editing. The observation of non-Mendelian segregation in two lines stimulated us to investigate if there is any new kind of base editing pattern in the T_1_ plants. We have sequenced the target locus amplified from 14 plants from *Z3*/10-105 and 12 plants from *Wsl5*/9-79.

Interestingly, we found that the 12^th^ base (A) of the *Z3* protospacer sequence has been changed to G in 7 plants out of 14 (Figure 3A). Out of these seven plants, two were homozygous mutants for the 12^th^ base, and five were heterozygous. The T_0_ plant 10-105 had exhibited editing only at the 7^th^ position of the protospacer (targeted). However, the editing of this 12^th^ A indicates the base editor is still active in the T_1_ generation. *pKABE7.10* showed an extended activity window in contrast to what reported in the mammalian cells (Gaudelli *et al.*, 2017).

Surprisingly, in the *Wsl5* line (9-79), two plants showed unusual base conversion in the activity window. One displayed a single G>A (4^th^ base) conversion, while the other showed a single A>T (5^th^ base) conversion (Figure 3B), that is an unexpected behavior of adenine base editor. The result provoked us to sequence more T_1_ plants for unraveling any unintended editing. We did not observe any other type of unwanted mutation in any of the lines. We then assumed if the active base editor could induce mutation in the other target locus in the single mutant plants in T_1,_ i.e., checking *Wsl5* locus in *Z3* mutant plant and vice versa. Unfortunately, no single plant was obtained with both the loci edited.

### Base conversion at the Wsl5 and Z3 loci translate to mutant phenotype

We further looked at whether the successful targeted base editing alters the phenotype of rice plants. A biallelic single nucleotide change (T>C) in the wild type *Zebra3* gene causes the mutant phenotype: transverse dark-green/green sectors in the mature leaves, late flowering and stunted plant growth (Kim *et al.*, 2018). The mutation in exon 3 causes a missense mutation from Serine542 to Proline542. Successful creation of the homozygous mutant (*z3/z3*) line by adenine base-editor displayed all the mutant phenotypes described by Kim et al. (2018) (Figure 4A-4D). As expected, stunted growth phenotype in heterozygous plants was not as severe as in homozygous plants (Figure 4B). We have noticed one or two of the leaves per mutant plant exhibited leaf variegation phenotype (Figure 4D), while others were normal. However, delayed flowering was common in all the *z3/z3* mutants. We observed 1-2 panicles per plant and fewer number of seeds per panicle in the homozygous mutant, while the panicle numbers in heterozygous (*Z3/z3*) plants were not significantly different from the wild type plants. We also observed a reduction in seed size in the mutant plants in comparison to the wild type Kitaake plants (Supplementary figure 2). Interestingly, the plants with the edited 12^th^ A in the *Z3* protospacer showed a similar phenotypic appearance as the *z3/z3* homozygous mutant (7^th^A in the protospacer) (Supplementary figure 3). The conversion of A>G at the 12^th^ position results in a missense mutation valine to alanine (V540A) (Fig 3A).

**Figure 4.**
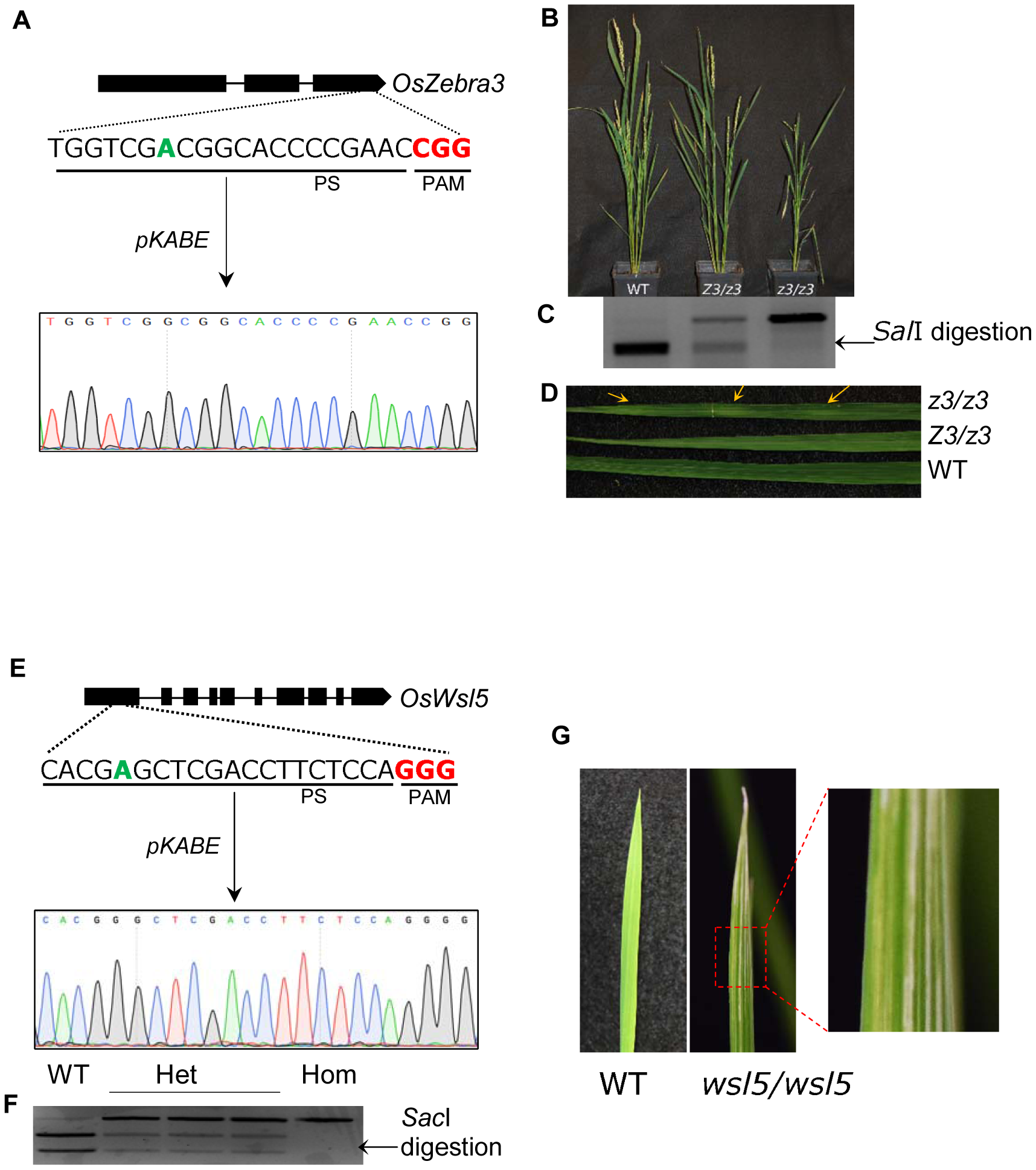
Characteristic phenotypes of *z3* and *wsl5* mutants generated by adenine base editors. **A.** Schematic representation of the target site in *OsZebra3 (Z3)* locus and DNA chromatogram of homozygous mutant (*z3/z3*) plant. **B.** Phenotypic appearance of wild type (WT), heterozygous (*Z3/z3*), and homozygous (*z3/z3*) mutant plants. **C.** Representative restriction analysis from WT, *Z3/z3*, and *z3/z3* plant showing complete, partial and nil digestion, respectively. **D.** Leaf variegation phenotype showing green and dark green sectors in mature leaf. Yellow arrow indicates green sectors. **E.** Schematic representation of the target site in *OsWsl5* locus and DNA chromatogram of homozygous mutant (*wsl5/wsl5*) plant. **F.** Representative RE analysis for WT, hetero-, and homozygous mutant plants. **G.** White strip leaf phenotype in homozygous (*wsl5/wsl5*) seedling.

The homozygous *wsl5* mutant is known to exhibit white-striped leaves in the seedlings (Liu *et al.*, 2018). A single nucleotide polymorphism (T>C) is the causal mutation of *wsl5*. It alters a CTC codon to CCC resulting in a Leucine151 to Proline151 amino acid substitution in the conserved region of the first exon of the *Wsl5* gene. We have noticed that the base-editing derived homozygous (*wsl5/wsl5*) mutant Kitaake seedlings exhibited white stripe leaf phenotype (Figure 4E-4G). Mature mutant plants displayed normal appearance and phenotype as the non-transformed wild type plants or transformed non-edited plants.

### Obtainment of transgene-free, base-edited mutants

Developing Cas9 or base editor-free, non-transgenic mutant plants is highly desirable to address the regulatory issues of genome edited crop plants. Segregation of integrated Cas9 transgene from the induced mutation in a distant locus would provide the possibility to readily obtain Cas9 free plants with the targeted mutation. While analyzing the segregation pattern of the mutants in T_1_ generation, we have obtained base-editor (nCas9-adenine deaminase) free *z3* and *wsl5* mutants. Among all T_1_ plants derived from both *wsl5* and *z3* monoallelic T_0_ mutant lines, we obtained 21% of plants devoid of the base-editor. The absence of the base editor was determined by the negative result in ABE (TadA) specific PCR (Supplementary figure 4). Both biallelic and monoallelic mutant plants were found to be transgene-free. What emerges from the results reported here is that we can generate precisely base edited plants free from integrated T-DNA within a very short period. Obtaining T_1_ plants with the desired edit and devoid of any foreign DNA has taken us only a single generation (about five to six months).

### Assessment of indels and off-target base editing

To further examine the specificity of the adenine base editor used in our study, we analyzed the base editing percentage in the potential off-target sites. Off-target of base editors might presumably occur at the potential genomic sites that could be targeted by Cas9 through partial homology with the guide RNA. According to CRISPR-GE (<u>skl.scau.edu.cn</u>) designated scores, off-target loci were selected. A total of six off-target loci, four for *Wsl5* and two for *Z3* were amplified using site-specific primers (Supplementary table 1) and sequenced. Genomic loci other than the two selected sites for *Z3* did not contain a targetable A in the activity window. Randomly selected 20 plants for each type of mutant was subjected to off-target analysis. No mutations including indels or single base substitutions was detected at any of these putative off-target sites (Table 2). Our data confirmed the high specificity and precision of adenine base editing in rice plants.

**Table 2.**
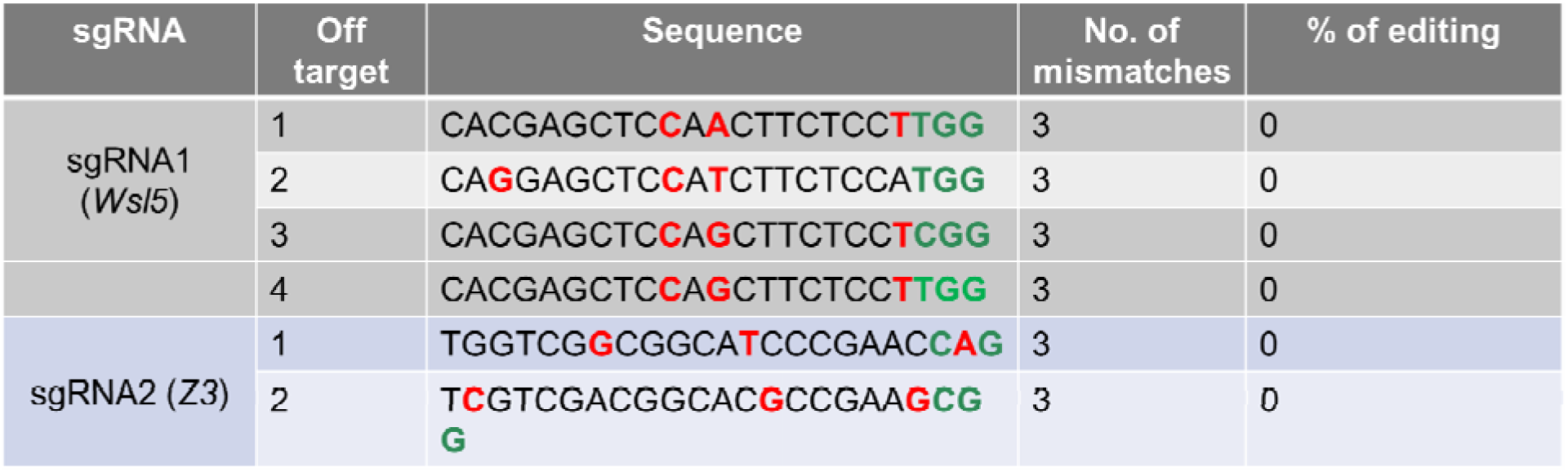
Off-target editing efficiency. Percent of editing includes both base conversion and indel generation.

## Discussion

In this study, we achieved successful and precise A>G base editing at two gene loci in the model plant, rice using two versions of adenine base editor, ABE7.10 and ABE7.9. The SpCas9 nickase (D10A) (nCas9) fused with deaminase (TadA-Tad* A) via a 32-amino acid {(SGGS)_2_-XTEN-(SGGS)_2_} linker was cloned under the control of a rice ubiquitin promoter for both transient expressions in protoplast and *Agrobacterium* meditated stable expression (Fig. 1A). The deaminase (TadA-TadA*) was fused at the N terminus of nCas9 as described in the original report (Gaudelli *et al.*, 2017). Fusion of adenosine deaminase at the C terminus has been demonstrated to be ineffective in rice and wheat (Li *et al.*, 2018). We used our synthetic polycistronic tRNA-gRNA (PTG) approach to efficiently produce two gRNAs for both the targeted genes (Fig 1B) (Xie *et al.*, 2015). The expression of PTG was driven by OsU3, a Pol-III promoter. As a tRNA gene contains the internal boxA and boxB which act as promoter elements for the RNA Pol-III, transcription of tRNA-gRNA is enhanced. After transcription as a unit, the endogenous RNase P and RNase Z would splice out the tRNA and as a result, individual sgRNA would be available to form complexes with the base-editor (nCas9-deaminase fusion). Earlier studies in rice reported adenine base editing with single guide RNA (Hua *et al.*, 2018*a*; Li *et al.*, 2018; Yan *et al.*, 2018). This system enables us to clone multiple gRNAs in a single vector to target multiple loci in the genome. In order to get a visible phenotype resulting from the base editing, we have targeted functional SNPs in two recently identified and cloned rice genes, viz., *white stripe leaf 5* (*Wsl5)* and *Zebra3 (Z3). Wsl5* encodes a pentatricopeptide repeat protein and a single nucleotide polymorphism (T>C) in the first exon caused Leucine_151_ to Proline_151_ substitution resulting the mutant *wsl5*. The *wsl5* mutant displays white-striped leaves at the seedling stage (Liu *et al.*, 2018). *Zebra3 (Z3)* is a putative citrate transporter gene and a single base substitution (T>C) in the third exon gives rise to a *z3* mutant plant which exhibits dark-green/green variegation in mature leaves (Kim *et al.*, 2018). In order to install T>C mutation in the coding strand, the opposite strand needs to be targeted by ABE. We designed sgRNAs targeting the opposite strand where target adenines were at protospacer position 5 and 7 for *Wsl5* and *Z3*gene, respectively.

For rapid verification of multiplexed base editing by ABE, we transfected rice protoplasts with the vector *pPr-ABE7.10* containing gRNA for both the targets. Target regions of *Wsl5* and *Z3* were amplified by PCR from the genomic DNA harvested from protoplasts at 3-4 days post transfection. As the target A bases are contained within recognition sites of *Sac*I and *Sal*I in *Wsl5* and *Z3* genes, respectively, successful base editing destroys the restriction sites. As evidenced from CAPS analysis, 24.14 % of mutated *Wsl5* was obtained, whereas the mutation frequency of *Z3* locus was about 13.31% (Fig. 1B). A recent study in rice and wheat protoplasts reported A>G conversion frequencies up to 7.5% (Li *et al.*, 2018). Kang et al. (2018) reported an A>G base editing frequencies up to 8.8% and 4.1% in Arabidopsis and rapeseed protoplast, respectively.

However, co-transfection of ABE plasmid and mutated GFP plasmid in rice protoplasts exhibited up to 32.8% of GFP fluorescent cells indicative of ABE mediated correction (Li *et al.*, 2018). When we sequenced the undigested products, we observed monoallelic targeted editing in both the loci (Fig.2). This finding, while preliminary, suggests that ABE works efficiently in rice. After this initial indication of successful A>G editing, we sought to generate stable transgenic rice plants harboring the intended mutation.

Since we observed a lower percentage of base editing at the *Z3* locus, we assumed the position of target A (7^th^ base of the protospacer) might be one of the influencing factors. Due to this assumption, we prepared two variants of adenine base editors *pKABE7.10* and *pKABE7.9*. Although *ABE7.10* was reported as the best one among the variants for the activity window ranging from the 4^th^ to 8^th^ base of the protospacer, ABE7.9 is better suited when the target base is at the 8^th^ to 10^th^ position. We reasoned that the window might be different for the plant system and 7^th^ is not favorable for ABE7.10. When we analyzed the regenerated plants from pKABE7.10, in contrast with the protoplast assay result, we obtained only 2.81% edited plants for *Wsl5* and 14% for *Z3*.

On the other hand, *pKABE7.9* performed far better for *Z3* with ∼39% editing, but poorer for *Wsl5* with only 1.34%. This very low editing efficiency at *Wsl5* locus could possibly be attributed to many reasons. When we were planning for the study, we had bioinformatically analyzed RNA fold pattern of sgRNA for both the loci and we noticed that *Wsl5* sgRNA was far better in its folding pattern with 3 stem-loop structures than the *Z3* sgRNA folding. This indicates the folding pattern is unlikely to have affected the editing outcome for *Wsl5* and *Z3* loci. The efficiency of Cas9 or base editors varies between different sgRNA target genes. For example, rice *PMS1* and *OMTN1* were reported to be resistant to base editing with the tested sgRNAs (Hua *et al.*, 2018*b*). Similarly, rice Tms9-1 gene was found to be resistant to ABE (Yan *et al.*, 2018). The low efficiency or nil base editing might be due to the poor accessibility of the locus to the base editor. Association with nucleosomes or other proteins may also reduce or hinder accessibility of the target base to the deaminase. Incidentally, *Wsl5* protospacer has a stretch of four G bases immediately after its 5’ terminus, which can act as three consecutive NGG PAMs and one NAG PAM. The presence of additional Gs next to PAM, which could shift the editing window, impede the rate of R-loop resolution or hinder Cas9’s alignment for proper gRNA–DNA interactions (Malina *et al.*, 2015). Proper R-loop resolution is the prerequisite for the generation of accessible single-strand DNA targeted by the base editor.

Although earlier studies showed ABEs applicability in rice base-editing, they have not reported the evidence of germline transmission of the mutation (Hua *et al.*, 2018*a*; Li *et al.*, 2018; Yan *et al.*, 2018). Analysis of segregation of monoallelic mutants in T_1_ generation revealed two lines from *Z3* and one line from *Wsl5* did not follow the Mendelian pattern. Most of the T_1_ plants from both the lines harbor the base editor transgene as evidenced by ABE specific PCR. Generation of new mutations in the T_1_ generation was likely due to the presence of active base editors. This kind of unpredicted segregation was reported in Cas9 treated *Arabidopsis* (Fauser *et al.*, 2014) and rice (Xu *et al.*, 2015; Ishizaki, 2016) plants. Plants descendent from mutants generated by active Cas9 are prone to further rounds of editing until the PAM and seed region of protospacer are destroyed by editing. However, the situation is quite different for the plants derived from base editing. If the activity window of base-editor contains only one targetable A, creation of biallelic germline mutation of the target in first generation would make the base editor unnoticeable in the following generation. Then the descendent from that mutant plant should all carry the homozygous mutation for that loci. However, if the generated mutation is monoallelic in the first generation, there are chances of activity of base editor in the next generation. That was likely the case in our study, which could explain why three of the tested lines did not follow Mendelian segregation pattern. The vigilant nature of base editors in the T1 generation was further evidenced by the detection of ‘out of window’ base editing (12^th^ base of the protospacer) by ABE7.10 in *Z3* mutant line (10-105) (Fig.3A). Earlier studies of ABEs in plants reported activity windows similar to that observed in mammalian cells (4 to 8 base position). However, Hua et al. (2018) reported editing at the 10^th^ base by SpCas9 nickase based-ABE7.10, and at 12^th^ and 14^th^ by SaCas9 nickase based-ABE7.10. It is evident that the activity window of a base editor varies with the target.

Unlike cytosine base editors (CBEs) which are reported to cause unintended C>A or C>G editing, ABEs are not known to generate undesired mutation other than the expected A>G (Reviewed by Molla K and Yang Y, 2019). Strikingly, two T1 plants exhibited unusual base conversion in our study (Fig.3B). In one plant, the target A was found to be converted to T. Mechanistically, ABE acts by deaminating deoxy-adenosine (dA) to deoxy-inosine (dI) which is read as guanosine by replication machinery, and as a result, it causes a post-replicative transition to G (Gaudelli *et al.*, 2017). Although deoxy-inosine:deoxy-cytidine (dI:dC) is the most stable pair, dI can also pair with dA (Alseth *et al.*, 2014). A dI:dA pairing would give rise to a post-replicative A>T conversion. That is one of the possible explanations for the A>T conversion we observed. In another plant, we observed that an adjacent G (6^th^base in the protospacer) was converted to A. Accidental deamination of guanine by tadA-tadA* may give rise to xanthine (X) which can subsequently pair with T (Herraiz and Galisteo, 2018). An X:T pair may result in the conversion of an original G-C base pair to A-T.

Most of the previous studies on rice have only reported evidence on genomic changes by ABE and not provided direct evidence of mutant phenotypes (Hua *et al.*, 2018*a*; Li *et al.*, 2018; Yan *et al.*, 2018). One of the reasons to select *Wsl5* and *Z3* as the target genes in our study to demonstrate ABE effectivity was that the base editing could be translated into a detectable phenotype. We have analyzed segregation patterns of the mutation and obtained mutant phenotypes in T_1_ plants. White striped leaf phenotype was evident in the *wsl5* mutant whereas z3 mutant exhibited altered phenotypes like transverse dark green/green sectors in mature leaf, shortened plant height, delayed flowering, and reduced panicle size in T_1_ plants (Fig.4). These results confirm the findings of the original studies on *Wsl5* and *Z3* genes (Kim *et al.*, 2018; Liu *et al.*, 2018).

In comparison to wild type plants, the *z3/z3* homozygous mutant in our study apparently exhibited far more reduced height than that reported by Kim et al. (2018). It is likely that this difference might be due to the use of different genotypic background in the present study (Kitaake) from that used in the earlier study (Kinmaze) (Kim *et al.*, 2018). Our study is the first following study on both the two genes providing data on the mutant phenotypes that validate their findings. Incidentally, as described earlier in the previous paragraph, we noticed editing of 12^th^ A in the protospacer of *Z3* which causes V540A mutation. We tracked the development of those plants carrying the heterozygous and homozygous mutation for 12^th^A and observed similar growth retardation. Homozygous plants showed a more prominent dwarf phenotype than the heterozygous plants (supplementary fig.2). This unexpected finding suggests that the change in 540^th^ amino acid can also cause a phenotype similar to the *z3* which was reported to occur due to alteration at 542^nd^ amino acids (Kim *et al.*, 2018). From this result, it can be assumed that the Valine540 is as critical as the S542 residue for the natural structure and function of Zebra3 protein.

Base editing allows us to rapidly and precisely introduce desired single nucleotide variation in the cultivated crop genome. In the present study, we developed targeted mutants for two rice genomic loci in less than one year. Generation of the same kind of genetic variation would have taken much longer time through traditional breeding techniques. Besides, genetic crosses including several rounds of backcrossing generate numerous nucleotide variants, often leads to undesirable effects as a result of genotype × genotype interactions (Huang *et al.*, 2016). By introducing SNPs, base editing can create an intended trait or attenuate an unwanted trait. Genetic modification (GM) technology, which depends on insertion, integration, and stability of DNA sequences from other species or the same species, faces tight regulation in many countries. Although base editing involves foreign DNA transformation, it does not rely on the permanent presence of that DNA sequence in the genome. After the creation of intended genetic mutation, base editing machinery is no longer needed. As edited plants without any foreign DNA remnants could address the issues of government regulation and public acceptance, generation of base editor free mutants is crucial. In rice, transgene-free mutants could be readily obtained through genetic segregation of the base editor. In the T_1_ generation, we obtained both homozygous and heterozygous *wsl5* or *z3* mutants which are Cas9-TadA-TadA* free. Earlier studies on plant ABE did not report the generation of base editor free mutants (Hua *et al.*, 2018*a*,*b*; Li *et al.*, 2018; Yan *et al.*, 2018). Since this base editor contains Cas9 nickase (nCas9) instead of fully active Cas9 and does not create a double-strand break (DSB), it minimizes the creation of DSB associated by-products such as indel, rearrangements, and translocation (Molla and Yang, 2019). We have not detected any indel generation in the targeted loci or in the studied potential off-target loci in the mutants. Our result corroborates with other findings in ABE induced mutant rice plants (Hua *et al.*, 2018*a*; Li *et al.*, 2018; Yan *et al.*, 2018). However, ABE induced indel generation (<0.1%) was reported in a transient assay using protoplasts (Kang *et al.*, 2018; Li *et al.*, 2018).

In contrast to ABE, cytosine base editors (CBEs) have been reported to generate indels up to 9.61% in rice (Li *et al.*, 2017) and up to 10-69% in tomato and rice (Shimatani *et al.*, 2017). The superior performance of ABEs in terms of generating very low indel mutation than CBEs might be due to the less active cellular inosine excision repair system than uracil excision repair system (Gaudelli *et al.*, 2017). Off-target editing by Cas9 or any other nucleases is one of the major concerns in genome editing experiments. Similarly, base editors can also generate off targets editing due to the nonspecific interaction of nCas9 with partially homologous loci. When we assayed editing in a total of six potential off-target loci, no off-target mutations were detected demonstrating the precise nature of ABE in rice. These results are consistent with the data obtained in two earlier studies in rice (Hua *et al.*, 2018*a*; Yan *et al.*, 2018). However, off-target editing may vary case-by-case depending on the unintended interaction of sgRNA and Cas9.

In conclusion, we have developed an adenine base editing system for plants and generated precise rice mutants for two distinct loci within a very short period of time by employing CRISPR/Cas-mediated adenine base editors. Our study demonstrates that ABEs can be used to generate precise and heritable base change mutations in plants and to rapidly develop edited, yet transgene-free, crops. This work will help pave the way to functionally validate many SNPs or achieve desired agronomic traits by installing A>G or T>C mutation in plant genome for natural and induced variants of genes.

## Methods

### Construction of base editing vectors

*pENTR11*-dual selection vector (Invitrogen) was digested with *Eco*RI and self-ligated to eliminate the *ccdB* gene. *pCMV-ABE7.10* (Addgene plasmid #102919) and *pCMV-ABE7.9* (Addgene Plasmid #102918) (Gaudelli *et al.*, 2017) were digested with *Not*I/*Pme*I to release the ABE-7.10 & 7.9 (TadA-Tad* A-nCas9) and cloned in the *Not*I/*Eco*RV site of *pENTR11*(-ccdB) to generate *pENTR11-ABE7.10* and *pENTR11-ABE7.9*, respectively. The *ABE7.10* was digested out from *pENTR11-ABE7.10* with *Bst*BI/*Xba*I and cloned in the same site of *pRGE32* (Xie *et al.*, 2015) replacing the *SpCas9* to generate *pPr-ABE7.10*.

Polycistronic tRNA-gRNA (PTG) for targeting both the rice *Wsl5* and *Zebra3* loci simultaneously was generated following our earlier described method (Xie *et al*., 2015). Primer sequences used for PTG synthesis are listed in supplementary table 1. The PTG fragment was digested with *Fok*I and inserted into the *Bsa*I digested *pPr-ABE7.10* vector. ABEs were expressed under the control of rice ubiquitin 10 promoter, while the PTG was transcribed under the control of OsU3 promoter. The *pPr-ABE7.10* was used for protoplast transfection.

For binary vector construction, the *pENTR11-ABE7.10* and *pENTR11-ABE7.9* was digested with *Bst*BI/*Xba*I to release *ABE7.10*and *ABE7.9*. The fragments were inserted into the same site of binary vector pRGEB32 (Xie *et al.*, 2015) replacing *SpCas9* to generate *pK-ABE7.10* and *pK-ABE7.9*. The PTG containing sgRNA for both *Wsl5* and *Zebra3* genes was inserted similarly into the *Bsa*I digested *pK-ABE7.10* and *pK-ABE7.9* vector.

### Protoplast Transfection

Rice (*Oryza stiva* ssp. *japonica* cv. Kitaake) protoplasts were isolated and transfected as described previously with few modifications (Xie and Yang, 2013). Briefly, rice stem and sheath were cut into 0.5-1 mm strips and immediately transferred into 10 ml of 0.6 M mannitol and incubated for 10 min. Mannitol was replaced with enzyme solution (1.5% Cellulase R10, 0.75% Macerozyme R10, 0.5 M mannitol, 10mM MES at pH 5.7, 1mM CaCl_2_, 5mM β-marcaptoethanol, and 0.1% BSA) and the rice tissues were digested for 5-8 hr in dark with gentle shaking. After adding 10-15 ml W5 solution (2mM MES at pH 5.7, 154 mM NaCl, 5 mM KCl, 125 mM CaCl_2_), protoplasts were filtered through 35 µm Nylon mesh. Protoplasts were pelleted down by centrifugation at 250g for 3 min and re-suspended in 1 ml W5 solution. After incubation for 1 hour at room temperature, W5 solution was removed by centrifugation and protoplasts were re-suspended in MMG solution (4 mM MES, 0.6 M Mannitol, 15 mM MgCl_2_) to a final concentration of 5×10^6^ cells ml^-1^.

PEG-mediated transfection was carried out using 200 μl of protoplasts and 20 μg of DNA (*pPr-ABE7.10*, and *pPr-ABE7.9*). A GFP expression cassette containing plasmid was used as the control to determine transformation efficiency. Protoplasts were transfected with ∼40% efficiency.

The protoplasts and DNA were gently mixed, and 1 ml of freshly prepared PEG solution (0.6 M Mannitol, 100 mM CaCl_2_, and 40% PEG4000) was added slowly. The mixture was incubated at room temperature for 20 min. Four ml of W5 solution was added to stop the transformation. Protoplasts were collected by centrifugation and resuspended in WI solution (4 mM MES, pH 5.7, 0.6 M Mannitol, 4 mM KCl) and transferred in six well plates. After 48 hrs of incubation, DNA was extracted from the protoplast for further analysis.

### Mutation detection by PCR, CAPS analysis and sequencing

The target regions for both *Wsl5* and *Zebra3* were amplified by PCR using specific pairs of oligo primers (Supplementary Table 1). DNA was extracted following a previously described method (Molla *et al.*, 2015). PCR products were digested with *Sac*I and *Sal*I for *Wsl5* and *Zebra3*, respectively. Since the target A for both the loci fall within a restriction enzyme recognition site, successful editing destroys the restriction sites. Digestion insensitivity would indicate base editing. After electrophoresis, the undigested DNA fragments were purified from agarose gels for DNA sequencing.

### Agrobacterium-mediated rice transformation

The base editing constructs were electroporated into *Agrobacterium tumefaciens* strain EHA105. The *Agrobacterium*-mediated transformation of 21-25 days old calli derived from Kitaake mature seed was performed as described (Mei *et al.*, 2007). Transformed calli were selected on hygromycin (50 µg/ml) containing media for 4-6 weeks and transferred to regeneration media. After regeneration of roots in rooting media, transgenic plantlets were transferred to soil and grown in controlled greenhouse for subsequent genotypic and phenotypic analysis.

### Analysis of base editing efficiency in the T_0_ generation

Genomic DNA was extracted from leaves of regenerated T_0_ plants as described previously (Molla *et al.*, 2015). The target loci were amplified by PCR using specific primers (Supplementary Table 1) and resulting DNA fragments were purified with PCR purification kit (Bio Basic Inc, Canada). The *wsl5* and *zebra3* PCR product was digested with *Sac*I and *Sal*I, respectively to determine the editing. The purified PCR products were also subjected to Sanger sequencing.

### Segregation analysis in the T_1_ generation and Phenotyping of mutant lines

Seeds obtained from monoallelic mutation of *Wsl5* and *Z3* were germinated to raise T1 plants. Ten plants from each line were evaluated by RE analysis and Sanger sequencing after targeted amplification. A chi-square test was performed to ascertain the inheritance pattern. T_0_ seeds from mutant plants were germinated and grown at 28°C/23°C (day/night) under 12/12 hr light and dark cycle. For phenotyping *wsl5* mutants, plants were grown at constant temperature of 20°C.

### Off-target analysis

Potential off targets of the base editing for both the genes were examined. The off-target sites having up to 3 base mismatches to sgRNA target region were identified using CRISPR-GE (http://skl.scau.edu.cn) software (Xie *et al.*, 2017). Then the corresponding sequence for Kitaake genome was retrieved from Phytozome and primers were designed using primer3 software (Untergasser *et al.*, 2012). A total of six, four for *wsl5* and two for *zebra3*, off-target loci were amplified, and analyzed with Sanger sequencing to detect any unwanted mutations.

## Acknowledgement

Kutubuddin Molla would like to acknowledge the United States-India Educational Foundation (USIEF), New Delhi and the U.S. Department of State for the Fulbright-Nehru Postdoctoral Fellowship. This work was supported by National Science Foundation Plant Genome Research Program Grant No. 1740874 and the USDA National Institute of Food and Agriculture and Hatch Appropriations under Project #PEN04659 and Accession #1016432 to Yinong Yang.

## Figure Legends

**Supplementary Figure 1.**
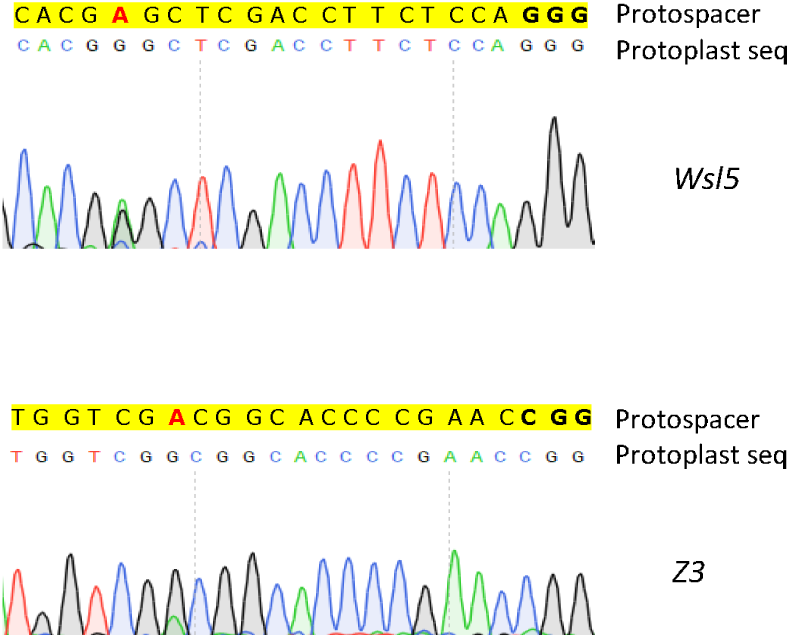
Sanger sequencing profile of targeted amplicon from protoplast DNA. In the protospacer seq, bold letters and red letter designate the protospacer adjacent motif (PAM) and target A base, respectively.

**Supplementary Figure 2.**
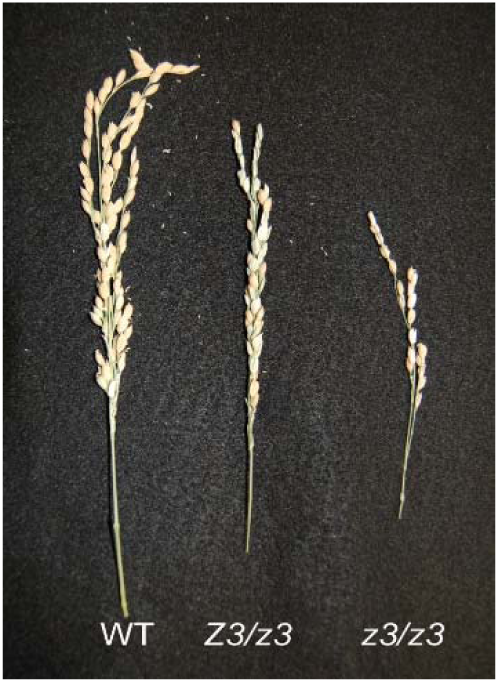
Panicles from wildtype (WT) and *z3* mutant plants.

**Supplementary Figure 3.**
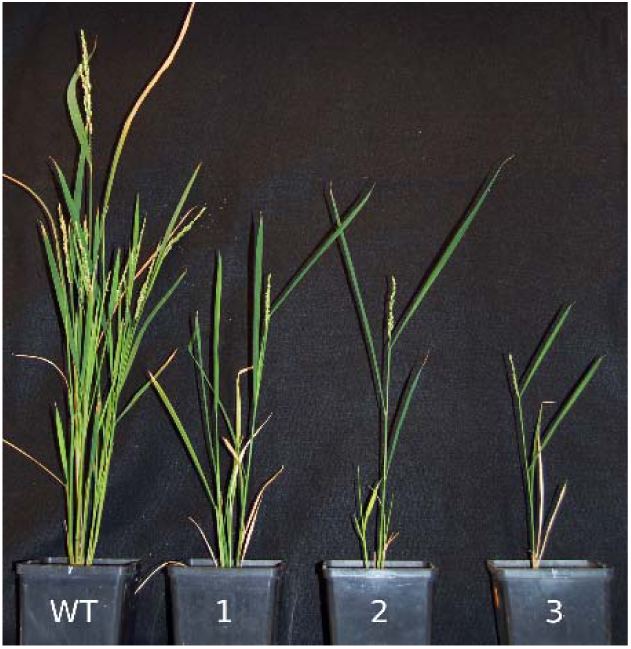
Phenotypic appearance of plants homozygous for 12^th^A→G in the *Z3* protospacer, which causes a Valine to Alanine missense mutation.

**Supplementary Figure 4.**
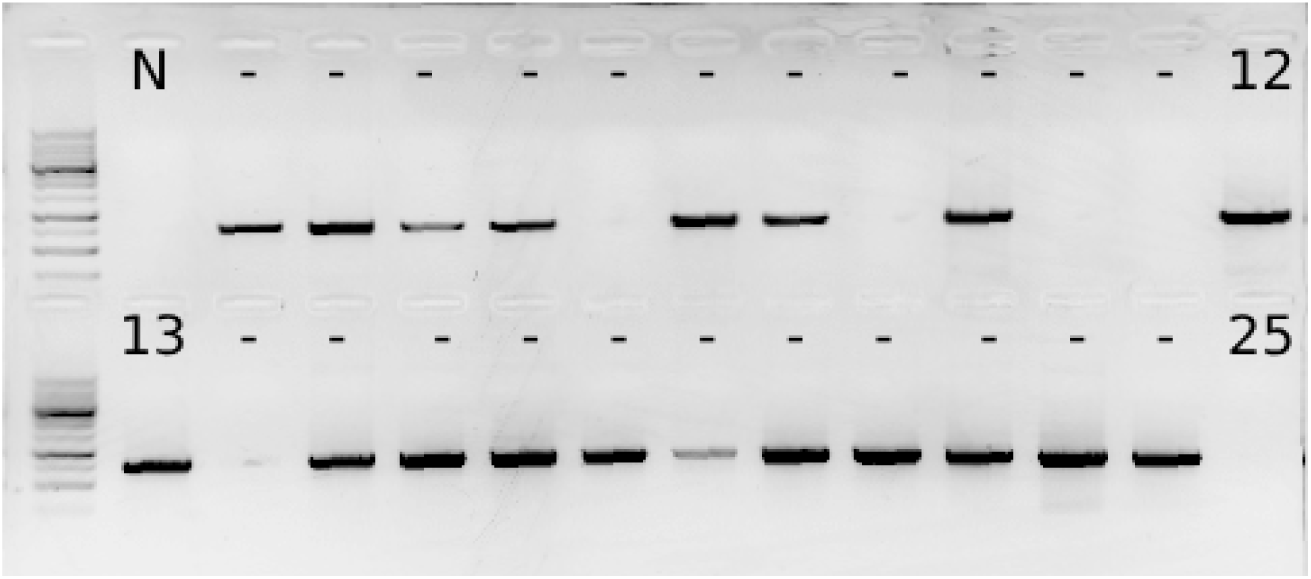
Representative gel image showing absence of base editors in the T1 mutants indicative of transgene free plants.

---

**Supplementary Table 1:**
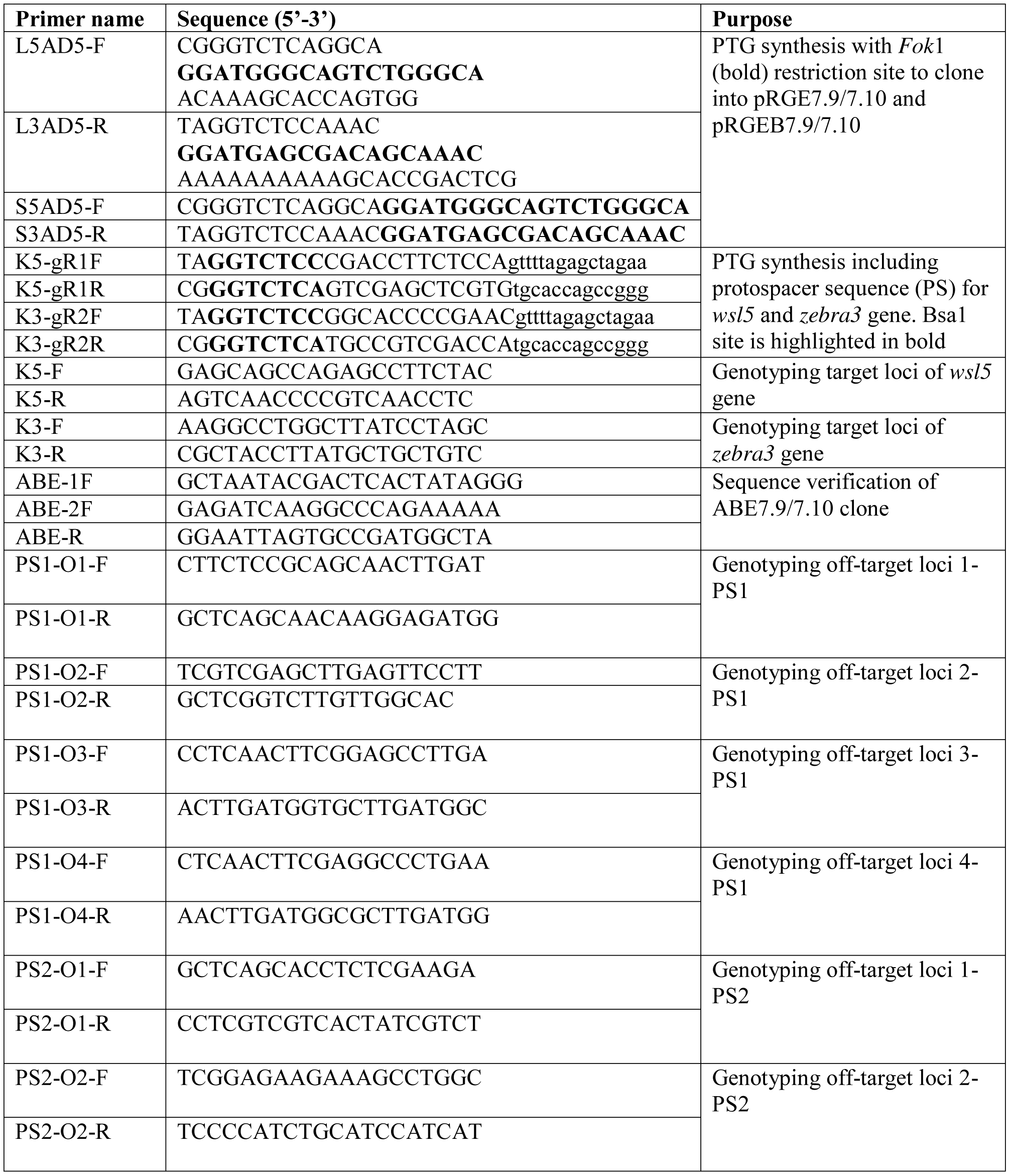
List of oligonucleotides ussed in the study (PS1-Protospacer for *Wsl5*, PS2-protospacer for *Zebra3*)

